## Supplementary figures and images for "Mammalian ZAP and KHNYN can independently restrict CpG-enriched avian viruses"

### Combined_ZC3HAV1_orthologues-PhyML_tree.pdf

Seaview Combined_ZC3HAV1_orthologues-PhyML_tree Thu Aug 21 10:02:51 2025

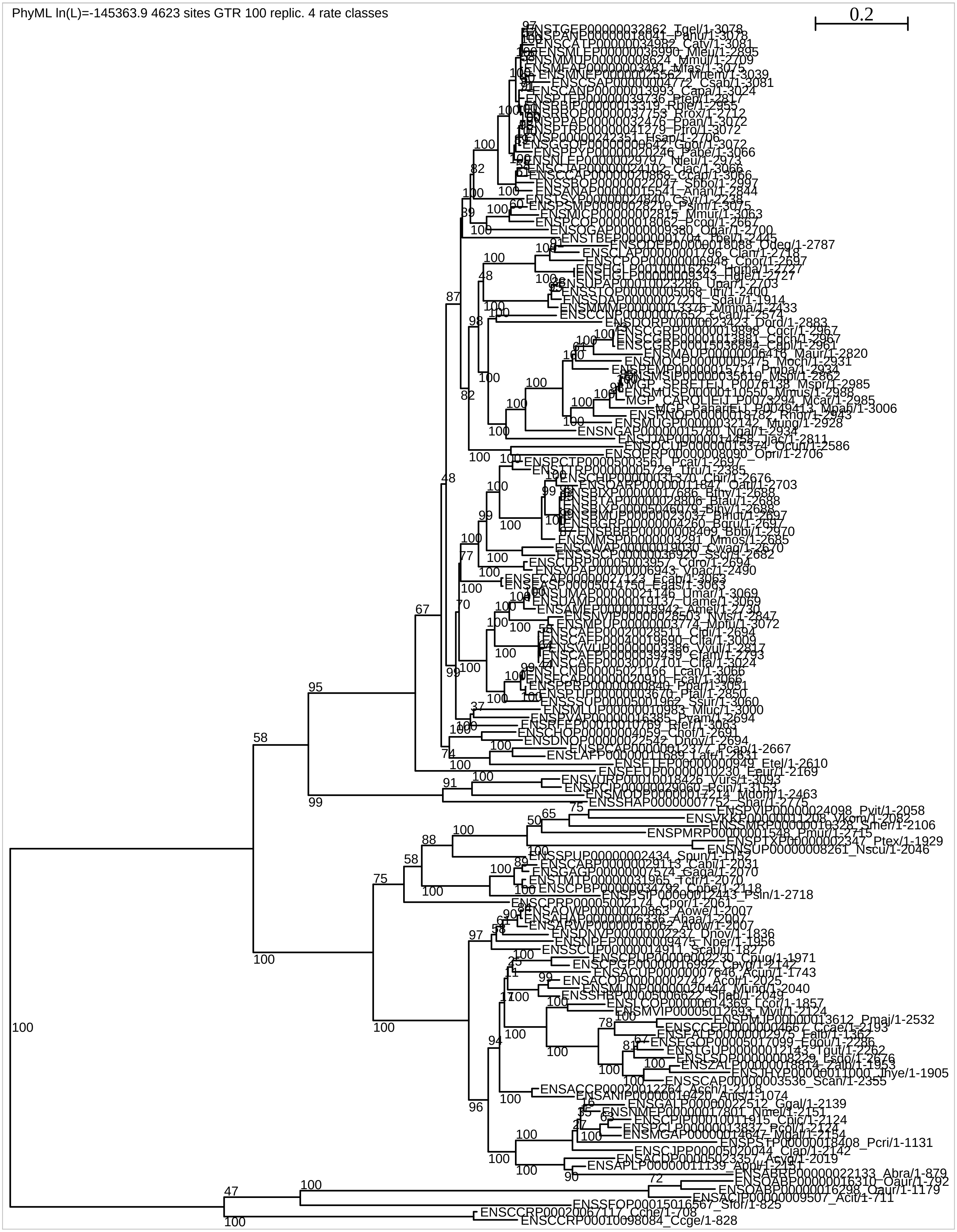

### FigS5C_PhyML_circular.pdf

Human+Zebrafish_KHNYN+hsaN4BP1_orthologues-PhyML_tree Thu Aug 5 12:22:47 2021 0.5

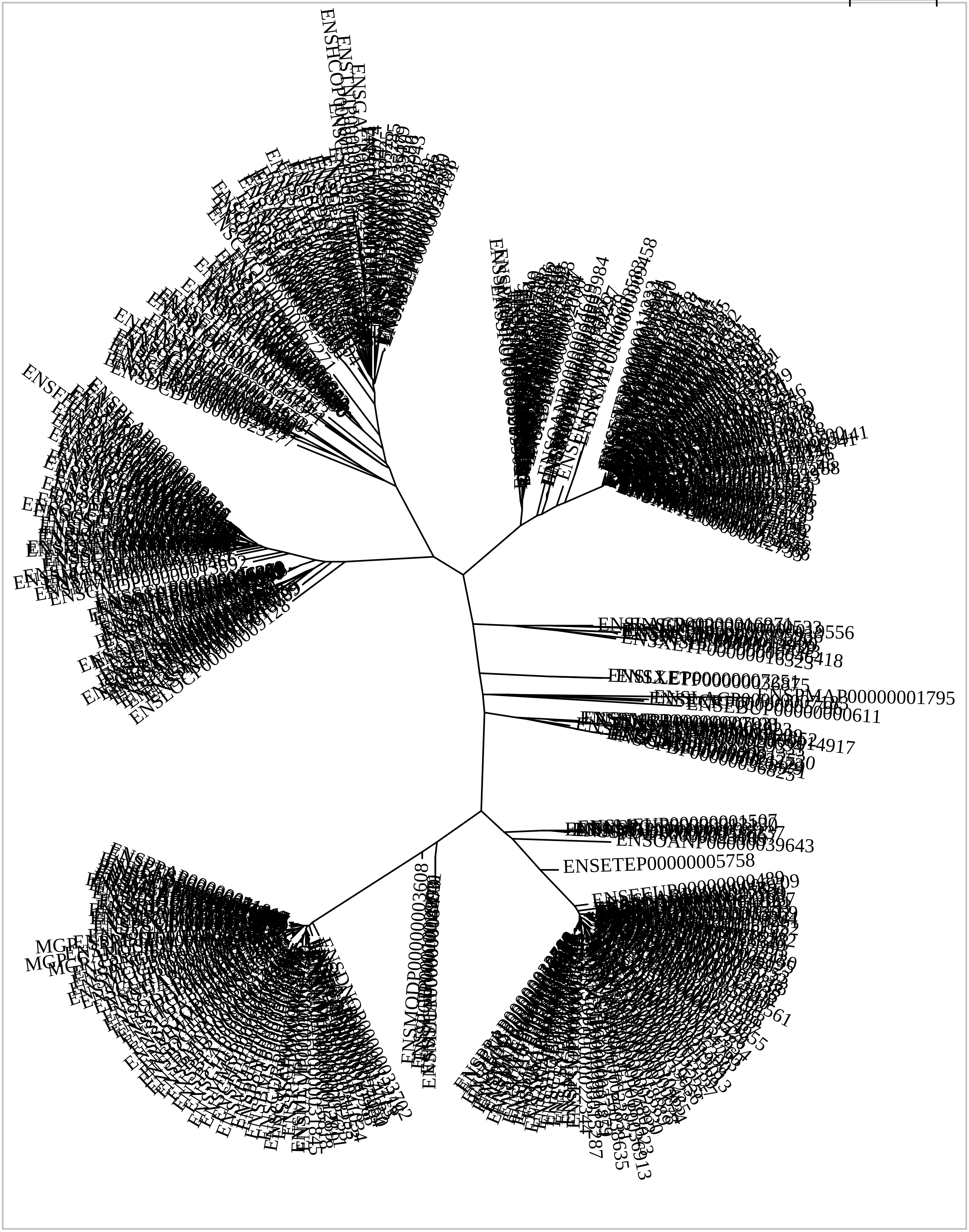
